# Cold exposure induces the constitutively active thermogenic receptor, GPR3, via ERRα and ERRγ

**DOI:** 10.1101/2025.08.09.669357

**Authors:** Olivia Sveidahl Johansen, Rebecca L. McIntyre, Janane Rahbani, Charlotte Scholtes, Damien Marc Lagarde, David Tandio, Astrid Linde Basse, Vincent Giguère, Lawrence Kazak, Zachary Gerhart-Hines

**Author notes:** Equal contribution (Olivia Sveidahl Johansen, Rebecca L. McIntyre).

## Abstract

**Objectives:** Despite transformative advances in obesity pharmacotherapy, safely increasing energy expenditure remains a key unmet need. Exploiting thermogenic adipocytes represents a promising target given their capacity for significant catabolic activity. We previously showed that G protein-coupled receptor 3 (GPR3) can drive energy expenditure in brown and white mouse and human adipocytes. GPR3 is a unique GPCR because it displays high intrinsic activity and leads to constitutive cAMP signaling upon reaching the cell surface. Therefore, the transcriptional induction of GPR3 is analogous to ligand-binding activation of most GPCRs. *Gpr3* expression is physiologically induced in thermogenic adipocytes by cold exposure and mimicking this event through overexpression in mice is fully sufficient to increase energy expenditure and counteract metabolic disease. Yet the factors mediating physiological *Gpr3* expression remain unknown.

**Methods:** Here, we apply ATAC-Seq to identify cold-induced promoter elements of *Gpr3*. We uncover a role for the estrogen-related receptors, ERRα and ERRγ, in the physiological transcriptional control of *Gpr3* using adipose-specific double knock-out mice with and without adeno associated virus (AAV)-mediated rescue.

**Results:** We show that ERRα directly binds the cold-induced promoter element of *Gpr3* and that adipocyte ERRα and ERRγ are required for the *in vivo* transcriptional induction of *Gpr3* during cold exposure. Importantly, deficient *Gpr3* cold-inducibility in adipose-specific ERRα and ERRγ KO mice is fully rescued by delivery of AAVs re-expressing either ERRα or ERRγ directly into brown adipose tissue.

**Conclusions:** ERRα and ERRγ are critical regulators of cold-induced transcription of *Gpr3* and represent a targetable strategy for pharmacologically unlocking GPR3-induced energy expenditure.

## 1. Introduction

Thermogenic adipocytes have a unique capacity for utilizing metabolites and lipids from the bloodstream to generate heat through catabolic processes [1–4]. These macronutrient-consuming and energy-dissipating activities are linked to cardiometabolic protection and enhanced glycemic control in humans [5–11], making thermogenic adipocyte activation an appealing strategy for treatment of metabolic disorders like obesity and diabetes. The activity of thermogenic adipocytes is highly orchestrated by G protein-coupled receptors (GPCRs) [12,13], with the Gα_s_-coupled β-adrenergic receptors serving as primary regulators of adipocyte catabolic recruitment. GPCRs represent the most druggable receptor class in biology [14–16], and therefore understanding the various GPCRs that control thermogenic adipose and their regulation is of significant interest for therapeutic purposes [13]. We and others recently discovered GPR3 as a potent activator of mouse and human thermogenic adipocytes [17,18]. GPR3 confers constitutive Gα_s_-coupled receptor signaling due to innate structural features of the receptor as well as a cell autonomously produced lipid ligand [17,19–21]. As a result, genetic or viral overexpression of GPR3 is alone sufficient to promote the thermogenic activity of adipocytes *in vitro* and *in vivo* and improve parameters of metabolic health [17]. Given that GPR3 begins increasing cAMP production as soon as it reaches the cell surface [22], the transcriptional control of *Gpr3* represents a key point of regulation. Therefore, transcriptional induction of GPR3 represents a stimulatory event comparable to the ligand-induced activation of more canonical GPCRs. We previously showed that environmental cold exposure acutely induces GPR3 transcription through a lipolytic signal [17]. Yet, the transcriptional machinery mediating this response remains unknown.

The estrogen-related receptors (ERRs) are a family of orphan nuclear receptor transcription factors that share sequence similarities with the estrogen receptors [23] and yet regulate distinct gene programs [24]. The three known subtypes of ERRs – ERRα, ERRβ, and ERRγ – are widely implicated in the control of cellular energy metabolism, including glucose and lipid homeostasis and mitochondrial biogenesis [25–27]. The ERRs are constitutively active [28,29]. However, protein coactivators, in particular peroxisome proliferator-activated receptor γ coactivator 1α (PGC1α) and peroxisome proliferator-activated receptor γ coactivator 1β (PGC1β), significantly impact their regulatory activity [27,29–31].

Critical roles of ERRα and ERRγ (together designated as ERRα/γ) in liver and muscle metabolism are well-established [27,32]. Moreover, recent advances have underscored the important roles of these nuclear receptors in the adrenergic-driven remodeling of brown adipose tissue (BAT) [33–35]. Whole-body ERRα KO mice are sensitive to cold exposure and display reduced BAT mitochondrial content [34]. The transcriptional response to β3-adrenergic stimulation in BAT was completely dependent on ERRα/γ and loss of adipocyte ERRα/γ impaired BAT thermogenic capacity [35]. The ERRs are more highly expressed in BAT than in white adipose tissue (WAT), with ERRα being the most prevalent, followed by ERRγ and then ERRβ [36,37]. Key coregulator *Ppargc1a* (encoding PGC1α) is also enriched in BAT and highly induced upon cold exposure [38]. Finally, the ERRs are proposed targets for activation in the treatment of metabolic diseases [27,35,39]. Recent optimization of a synthetic pan-ERR agonist, SLU-PP-332, with pharmacokinetic properties that allowed *in vivo* testing suggested that targeting ERRs may be a pharmacologically viable strategy [40].

In the present study, we investigate the chromatin dynamics surrounding the *Gpr3* transcriptional start site (TSS) in response to cold exposure, when *Gpr3* is transcriptionally activated. We identify a cold-responsive promoter element of *Gpr3*, show that ERRα is enriched at the promoter, and establish that ERRα/γ are required for physiological *Gpr3* cold induction through viral targeted rescue in tissue specific KO mice.

## 2. Materials & Methods

### 2.1 293T cell culture

293T cells (ATCC, 293T) were cultured in DMEM (Wisent, 319-005-CL) supplemented with 10% FBS (Multicell, 098150) and 1% penicillin/streptomycin (Multicell, 450-201-EL). The cells were maintained at 37 °C in a humidified atmosphere with 5% CO_2_.

### 2.2 Animals

The mouse studies performed adhered to the guidelines established by the Canadian Council of Animal Care and were approved by the Animal Resource Centre at McGill University. The mice were kept on a 12-hour light and 12-hour dark schedule with lights on at 07:00 h. The mice had access to a low-fat diet (2920X, Envigo) and drinking water *ad libitum*. They were maintained in an enriched environment with bedding and shredded paper strips and housed in groups of 3-5 per cage at 22 °C ± 2 °C, until they were 7-12 weeks old, when experiments were initiated. During the 30 °C acclimation and cold exposure experiments, the mice were housed in separate cages with bedding and shredded paper strips. Experiments were conducted on age-matched littermates at the specified temperatures. The mice were euthanized by cervical dislocation, and interscapular BATs were snap-frozen in liquid nitrogen and preserved at −80 °C for subsequent analysis. AdipoQ-Cre *Esrra/g* KO mice were obtained by breeding AdipoQ-Cre (B6.FVB-Tg(AdipoQ-Cre)1Evdr/J: JAX stock #028020 [41]) mice maintained on a C57BL/6J background to ERRα/γ^fl/fl^ mice [35,42–44]. The sexes of the mice used for experiments are indicated in the figure legends. Genotyping primer sequences are available in Supplementary Table 3.

### 2.3 Power calculation for AAV-mediated rescue study

The primary experimental output was *Gpr3* induction during 24 h cold exposure. Therefore, group size was determined using R, based on *Gpr3* expression in BAT following a pilot 24 h cold exposure study.

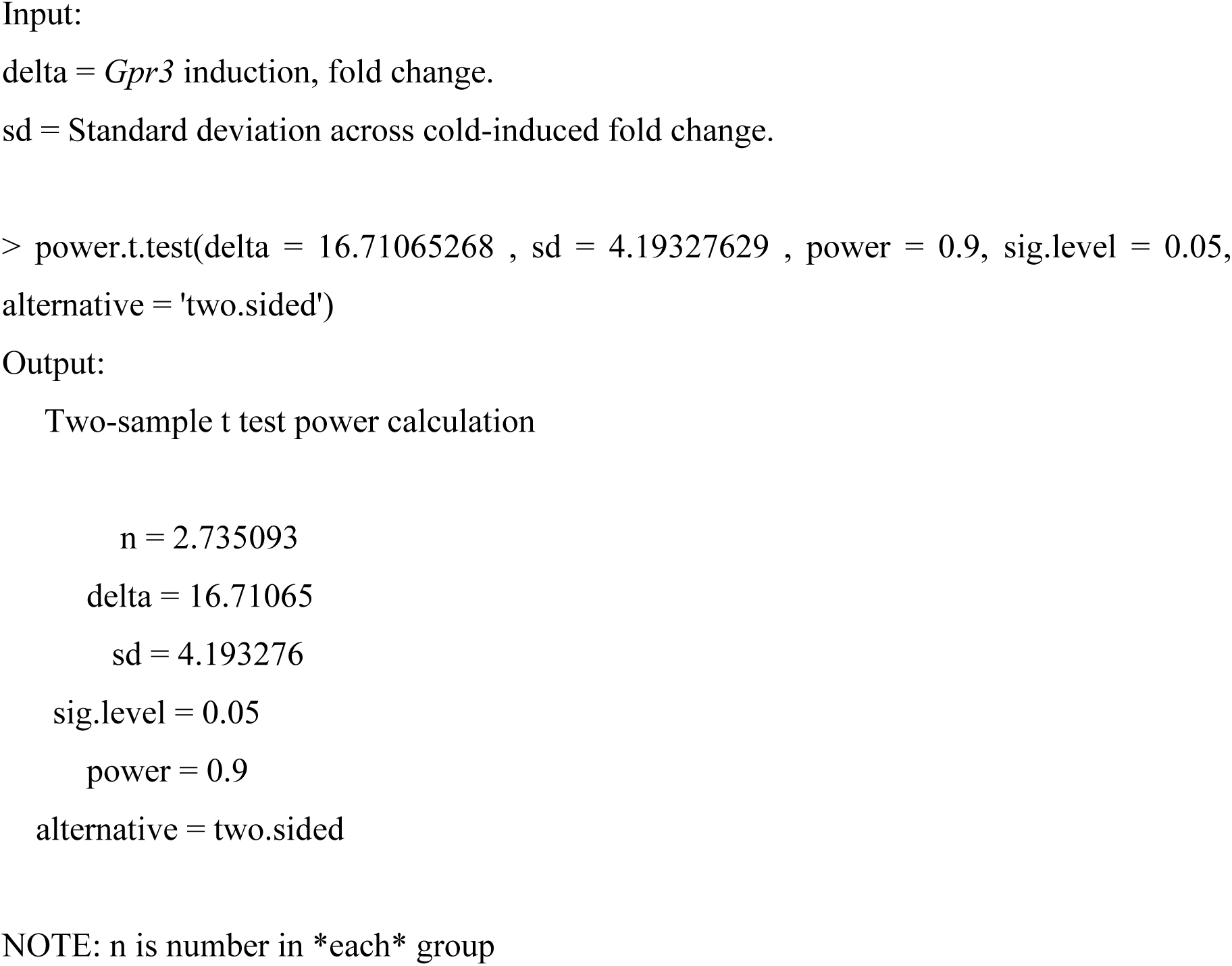

Male and female mice were included in the study. Experiments with male and female mice were conducted independently. 2-3 mice of each sex were included. No sex-dependent differences were observed. Data are pooled in the figures.

### 2.4 Site-directed adeno-associated virus (AAV) delivery

The AAV backbone for generation of AAV plasmids was kindly provided by the lab of Professor Christian Wolfrum (ETH Zurich). 293T cells (ATCC) were seeded at 80-90% confluence in a 15-cm cell culture plate. Immediately before transfection, media was replaced with 20 ml DMEM (Wisent, 319-005-CL) supplemented with 10% FBS (Multicell, 098150) and 1% penicillin/streptomycin (Multicell, 450-201-EL). 10 μg of targeting vector plasmid and 39.5 μg of helper plasmid pDP8 (Plasmid Factory, PF421-180518) were mixed in 2.5 ml Optimem (Gibco, 31985) before addition of 200 μl PEI (1mg/ml) (Polysciences, 23966-1). The transfection mix was briefly vortexed and incubated for 10 min. at RT. Then, the transfection mix was added dropwise onto the cells. The culture medium was refreshed 24 h after the transfection. Culture medium was then collected 72 h after transfection, and virus was concentrated using AAVanced Concentration Reagent (System Biosciences, AAV100A-1) in accordance with the manufacturer’s guidelines.

Site-directed delivery of AAVs into BAT was carried out as previously described [45]. The mice were anaesthetized by isoflurane at a concentration of 2.5% for induction and 1.5% for maintenance, and local (Lurocaine, Vetoquinol and Bupivacaine injection BP, Sensorcaine in 1:1 ratio) and general (Carprofen, 5 mg per kg body weight) (Rimadyl injectable solution, Zoetis Canada) analgesia were administered. Body temperature was maintained using a heating pad. To expose the brown fat depot, a longitudinal incision of 0.5-1.0 cm was made at the interscapular region. Using a 10 ml Hamilton syringe, 20 μl of AAV (≈10^12^ vg/ml) were administered across multiple (at least 5) injections within each lobe of the interscapular brown fat depot. This approach was employed to achieve a more homogeneous and extensive viral spread throughout the tissue. The incisions were closed by surgical clips and surgical glue (3M Vetbond^TM/TC^). The mice were monitored daily and received general analgesia by subcutaneous injections on days 1 and 2 post-surgery. 9 days post-surgery, the mice were single-housed, and the surgical clips were removed.

### 2.5 Cold exposure

The mice were allowed 5 days to acclimate to 30 °C before the temperature challenges. The specific duration and temperatures of the cold exposures are stated in the figure legends.

### 2.6 RNA extraction

RNA was extracted from snap-frozen interscapular BAT samples using QIAzol (Qiagen, 79306) and purified with RNeasy Mini spin columns (Qiagen, 74104) according to the manufacturer’s instructions. RNA was quantified using a NanoDrop 8000 Spectrophotometer (Thermo Scientific Pierce) prior to cDNA synthesis.

### 2.7 RT–qPCR

RNA was reverse transcribed using a High-Capacity cDNA Reverse Transcription kit (Applied Biosystems, 4368813). 10 ng cDNA and 150 nmol of each primer were mixed with 3 μl GoTaq qPCR Master Mix (Promega, A6001) to a final volume of 6 μl per sample. RT– qPCRs were performed using a CFX384 real-time PCR system (Bio-Rad) in a 384-well format, using the following cycling conditions: one step at 95 °C for 3 min., then 95 °C for 10 s, 60 °C for 20 s, 72 °C for 10 s. Expression data were quantified by ΔCt calculations normalized to housekeeping genes *36b4* (*Rplp0*) or *Ppib*.

### 2.8 Western blots

Samples were homogenized in lysis buffer (50 mM Tris, pH 7.4, 500 mM NaCl, 1% NP40, 20% glycerol, 5 mM EDTA, and 1 mM PMSF), supplemented with Roche protease inhibitors (Roche, 11836170001) using the TissueLyser II for 10 min. at 20 Hz. The homogenates were cleared by centrifugation at 20,000 g for 10 min. at 4 °C. Protein concentrations were determined using a bicinchoninic acid assay (Pierce, 23225) according to the manufacturer’s protocol. Protein lysates were denatured in Laemmli buffer (Bio-Rad, 161-0747) supplemented with 10% β-mercaptoethanol and incubated at 95 °C for 10 min. The samples were resolved by 10% Tris/Glycine SDS–PAGE and transferred to a polyvinylidene difluoride membrane. Following the transfer, the membranes were blocked for 1 h in TBS containing 0.05% tween (TBS-T) and 5% milk powder. The membranes were washed 3 times for 10 min. in TBS-T prior to incubation with primary antibodies (antibody dilutions are stated in supplementary table 2). Primary antibodies were diluted in TBS-T containing 5% BSA and 0.02% sodium azide. The membranes were incubated with primary antibodies at 4 °C overnight. On the following day, the membranes were washed 3 times for 10 min. in TBS-T prior to incubation with secondary antibodies. Secondary antibodies were diluted in TBS-T containing 5% milk powder. Membranes were washed 3 times for 10 min. before visualization with enhanced chemiluminescence western blotting substrates (Bio-Rad, 1705060).

### 2.9 ChIP-qPCR

ChIP-qPCR was performed as previously described [44]. Nuclei were isolated from 3 interscapular BAT pads per condition, each from individual WT mice. BAT pads were stroked in a nuclei preparation buffer (10 mM HEPES pH7.5, 10 mM KCl, 1.5 mM MgCl2, and 0.1% NP40), 25X with pestle A and 15X with pestle B, and the solution was filtered through a 100 μm strainer. Next, the nuclei were fixed with formaldehyde (1% final) for 12 min. at RT, quenched by 125 mM of glycine for 10 min., and washed twice with PBS/0.1% NP40. Subsequently, chromatin was sonicated in 1 ml of sonication buffer (50 mM Tris-HCl pH 8.1, 10 mM EDTA, and 1% SDS) to obtain fragments of around 500 bp. 20 μg chromatin DNA was diluted in ChIP dilution buffer (16.7 mM Tris-HCl pH 8.1, 1.1% Triton X-100, 167 mM NaCl, 1.2 mM EDTA, and 0.01% SDS) up to 2 ml. 10 μl of anti-ERRα antibody (Abcam, ab76228) were added to the sonicated chromatin and left to rotate overnight at 4 °C. The next day, 50 μl of Dynabeads protein G (Thermo Fisher Scientific, 10009D) were washed twice with PBS containing 0.5% tween and 0.5% BSA and added to the chromatin for 1 h under rotation at 4 °C. Next, beads were washed twice with 1 ml of cold low salt RIPA buffer (20 mM Tris-HCl pH 8.1, 0.1% SDS, 1% Triton x-100, 1 mM EDTA, 140 mM NaCl, and 0.1% Na-deoxylcholate), twice with 1 ml of cold high salt RIPA Buffer (20 mM Tris-HCl pH 8.1, 0.1% SDS, 1% Triton x-100, 1 mM EDTA, 500 mM NaCl, and 0.1% Na-deoxylcholate), twice with 1 ml of cold LiCl wash buffer (10 mM Tris-HCl pH 8.1, 250 mM LiCl, 0.5% NP40, 0.5% Na Deoxycholate, and 1 mM EDTA), and twice with RT TE buffer (10 mM Tris-HCl pH 8.0 and 1 mM EDTA). DNA was eluted overnight at 65 °C with 100 μl of ChIP elution buffer (10 mM Tris-HCl pH 8.0, 5 mM EDTA, 300 mM NaCl, 0.1% SDS, and 5 mM DTT) and 16 μl reverse cross-linking mix (250 mM Tris-HCl pH 6.5, 62.5 mM EDTA pH 8.0, 1.25 M NaCl, 5 mg/ml Proteinase K, and 62.5 ug/ml RNAse A) were added. Finally, chromatin immunoprecipitated DNA was purified using a QIAquick PCR purification kit and eluted in 31 μl of elution buffer (10 mM Tris-HCl pH 8.0 and 0.1 mM EDTA). Relative ChIP fold enrichments were controlled by inputs and normalized to the average of two non-specific control regions using a LightCycler 480 (Roche) and SYBR Green I Master Mix (Roche, 4887352001) as previously published [44]. Gene-specific primers used for ChIP-qPCR analysis are listed in supplementary table 3. Results represent the average of 3 biological replicates.

### 2.10 ATAC-Seq

Data was extracted from a published dataset [44] and analyzed as previously described [44].

### 2.11 Statistical analyses

Statistical analyses were performed using GraphPad Prism 9. Applied statistical analyses are stated in the figure legends.

## 3. Results

Adipose *Gpr3* is transcriptionally activated in response to cold exposure [17]. Therefore, we hypothesized that a cold-sensitive transcriptional element may be located within or near the *Gpr3* locus. We reanalyzed a dataset recently published by Rahbani *et al.* in which transposase-accessible chromatin sequencing (ATAC-Seq) was used to examine BAT nuclei isolated from mice housed at thermoneutrality (30 °C) or exposed to a 24 h cold challenge (6 °C) [44]. Here, we identified a differentially accessible region immediately upstream of the *Gpr3* TSS (peak position −121 relative to *Gpr3* TSS, GRCm38/mm10), indicative of a proximal promoter (**Fig. 1A**). The promoter element contained three estrogen related receptor elements (ERREs) (**Fig. 1B**), in line with the notion that the ERRs preferably bind in close proximity to the promoters of genes [27]. The ERRs are enriched in BAT compared to WAT [36,37] and directly regulate expression of genes important for oxidative and thermogenic processes [27,34,35,37,46–48]. Collectively, this prompted us to explore a potential role of the ERRs in the transcriptional regulation of *Gpr3*. Because ERRα is more highly expressed in BAT than the other ERR isoforms [36], we assessed ERRα-binding at the promoter region. We performed using chromatin immunoprecipitation coupled to quantitative PCR (ChIP–qPCR), testing two sets of primers, each covering the identified ERREs. ERRα was significantly enriched at the promoter compared to a control region that does not bind ERRα (**Fig. 1C**). Of note, the binding was independent of housing temperature (**Fig. 1C**). Thus, cold exposure activates a proximal promoter of *Gpr3*, which is enriched for ERREs that bind ERRα.

**Figure 1:**
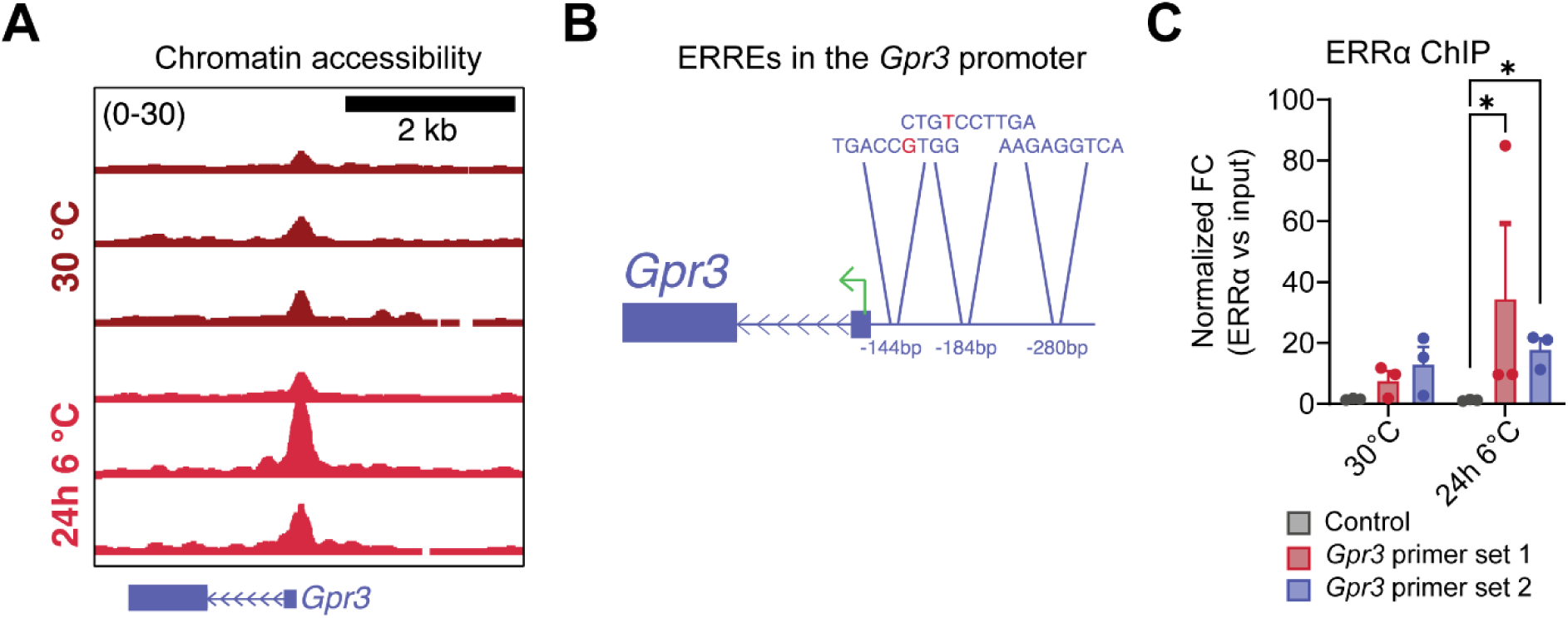
Identification of ERRα/γ as potential regulators of *Gpr3* cold induction. A. ATAC-Seq tracks showing cold-induced differentially expressed region (DAR) upstream of *Gpr3*. BAT nuclei were isolated from mice housed at 30 °C and following 24 h cold exposure at 6 °C (n = 3 per group, males). Data obtained from Rahbani *et al.* [44], B. ERREs located in the identified DAR. Red letters indicate nucleotides aberrant from known ERRE variants [24,60–63], C. ChIP-qPCR of ERRα bound to *Gpr3* DAR. BAT nuclei were isolated from mice treated as in A. Two different primer sets covering the DAR were tested. Data were log_10_ transformed prior to analysis by two-way analysis of variance (ANOVA); post hoc comparison using Šídák’s multiple comparisons test. Data are shown as the mean + SEM; *P ≤ 0.05,

We next explored whether the ERRs regulate *Gpr3* transcription *in vivo*. Because ERRγ (encoded by *Esrrg*) can compensate loss of ERRα (encoded by *Esrra*) in BAT [35], an adipocyte-selective *Esrra* and *Esrrg* double knock-out mouse model was selected for this study (hereby referred to as AdipoQ-Cre *Esrra/g* KO) (**Fig. 2A-B**). Adipocyte-specific ablation of both *Esrra* and *Esrrg* significantly blunted the cold induction of *Gpr3* (**Fig. 2C**), supporting the hypothesis that these nuclear receptors play a central role in the physiological control of GPR3.

**Figure 2:**
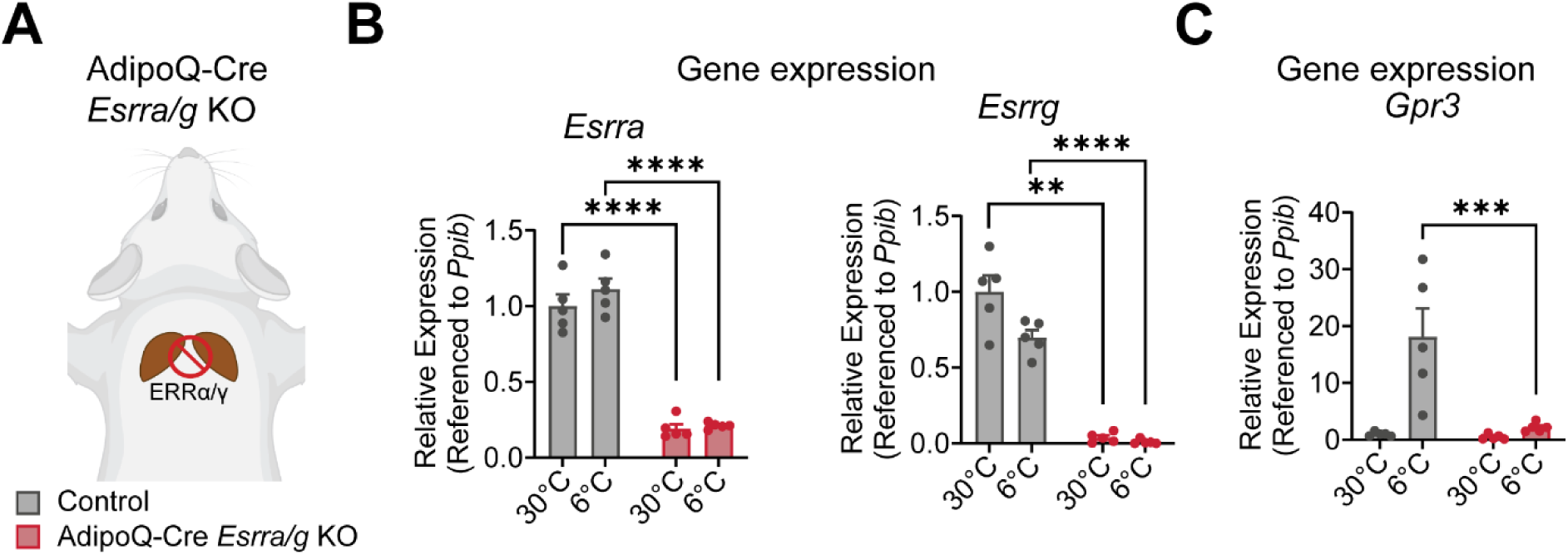
ERRα/γ are required for *Gpr3* cold-inducibility *in vivo*. A. Illustration of AdipoQ-Cre *Esrra/g* KO mouse model, B. RT-qPCR measuring gene expression of *Esrra* and *Esrrg* from BAT isolated from mice housed at 30 °C and following 24 h cold exposure at 6 °C (n = 5 per group, females). Data obtained from Rahbani *et al.* [44], C. RT–qPCR measuring gene expression of *Gpr3* from BAT isolated from mice treated as in B. Data presented as fold change. RT–qPCR data were log_10_ transformed prior to analysis by two-way ANOVA; post hoc comparison, testing genotype effect at each temperature using Šídák’s multiple comparisons test. Data are shown as the mean + SEM; **P ≤ 0.01, ***P ≤ 0.001. ****P ≤ 0.0001.

Mice lacking ERRα/γ display compromised BAT remodeling in response to pharmacological β3-adrenergic agonism [35]. Therefore, the effects we observe on *Gpr3* could be secondary to diminished adipocyte function. To determine if the impaired *Gpr3*-inducibility in this mouse model was directly a result of ERRα/γ loss-of-function, we established an acute gene rescue protocol. We restored ERRα or ERRγ expression in the interscapular BAT depots of the *Esrra/g* KO mice using site-directed delivery of AAVs expressing *Gfp*, *Esrra*, or *Esrrg* under the control of an uncoupling protein 1 (*Ucp1*) promoter (**Fig. 3A**). *Ucp1* is uniquely expressed in thermogenic adipocytes [1]. Thus, the promoter, in combination with site-directed delivery of AAVs enabled BAT-targeted ERRα/γ rescue. Two weeks after viral delivery, the mice were exposed to a 24 h cold challenge (4 °C), and BAT depots were isolated for analysis. The rescue was confirmed by qPCR and western blot (**Fig. 3B-C**). In fact, AAV-mediated gene transfer resulted in higher transcriptional expression of both *Esrra* and *Esrrg* compared to WT controls during cold exposure (**Fig. 3B**). This is likely due to the temperature-sensitive nature of the *Ucp1*-promoter driving the AAV-mediated overexpression. ERRα was only rescued in the cold exposed group (**Fig. 3C**).

**Figure 3:**
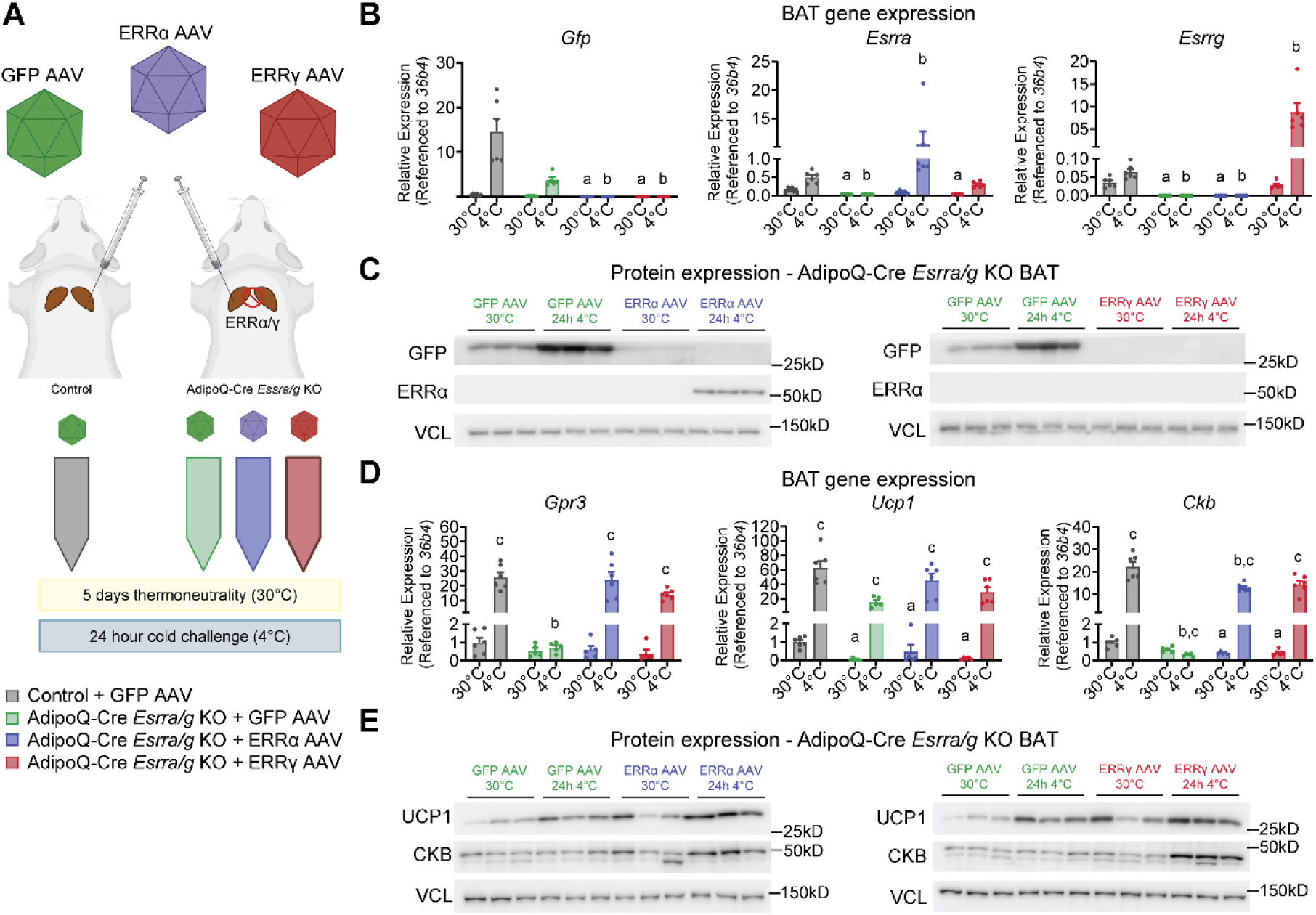
AAV-mediated rescue of *Gpr3* induction in AdipoQ-Cre *Esrra/g* KO mice. A. Illustration of AAV-mediated rescue protocol. Site-directed delivery of AAVs expressing *Gfp*, *Esrra*, or *Esrrg* was performed in control and AdipoQ-Cre *Esrra/g* KO mice. After recovery, the mice were housed at 30 °C or challenged with 24 h cold exposure at 4 °C. B. RT–qPCR from BAT isolated from mice treated as in A (n = 5-6 per group, females and males. Data points are pooled from two independent experiments). RT–qPCR data were log_10_ transformed prior to analysis by two-way ANOVA; post hoc comparison between control groups and AdipoQ-Cre *Esrra/g* KO groups using Šídák’s multiple comparisons test. a=at 30 °C, significantly different from control group. b=at 4 °C, significantly different from control group. C. Western blot from BAT isolated from mice treated as in A (n=3 per group, males). D. RT–qPCR from BAT isolated from mice treated as in A. RT–qPCR data were log_10_ transformed prior to analysis by two-way ANOVA; post hoc comparison between control groups and AdipoQ-Cre *Esrra/g* KO groups and between temperatures within each genotype using Šídák’s multiple comparisons test. a=at 30 °C, significantly different from control group. b=at 4 °C, significantly different from control group. c=significantly different between 30 °C and 4 °C within group. Data presented as fold change. (n = 5-6 per group, females and males. E. Western blot from BAT isolated from mice treated as in B (n=3 per group, males).

We assessed gene and protein expression of *Ucp1*/UCP1 to compare the transcriptional regulation of *Gpr3* to another cold-induced ERRα/γ target [35]. While the levels of *Ucp1* were reduced in the AdipoQ-Cre *Esrra/g* KO mice at either temperature, the cold-inducibility of *Ucp1*/UCP1 was not dependent on the ERRs (**Fig. 3D-E**). Our observations are in line with previous cold exposure studies of ERRα whole-body KO mice [33,34].

Strikingly, rescue of either ERRα or ERRγ restored cold-inducibility of *Gpr3* transcription in AdipoQ-Cre *Esrra/g* KO mice (**Fig. 3D**). As with *Gpr3*, it has previously been established that *Ckb* is transcriptionally cold-induced by the ERRs [44]. Additionally, BAT-specific GPR3 overexpression itself enhances *Ckb* mRNA levels [44]. Since there is currently no GPR3 antibody available, we used CKB as a proxy to validate the functional rescue on protein level. Cold-inducibility of CKB was rescued by AAV-mediated delivery of either ERRα or ERRγ (**Fig. 3E**). Interestingly, the CKB rescue was more pronounced in the mice that had been infected with ERRγ AAVs, which could be explained by a less coactivator-dependent activity of this ERR isoform [37,49,50].

## 4. Discussion

The ERRs are orphan nuclear receptor transcription factors with no known endogenous ligand. The ligand binding pocket of ERRα was initially thought to be very restricted based on a crystal structure of the ligand binding domain complexed with PGC1α [29]. However, Kallen *et al.* uncovered the potential for multiple dramatic conformational changes in the ligand binding pocket of ERRα, which supports the possibility of a bulkier endogenous ligand [51]. Intriguingly, activity studies of ERRα *in vitro* showed that the constitutive activity of the receptor was dependent on a serum-derived factor that was withdrawn upon charcoal treatment [52]. It is tempting to speculate that a lipolysis-regulated metabolite would provide the missing link between lipolysis and the role of these receptors in the transcriptional control of *Gpr3*. Of note, the major adipocyte lipase, hormone sensitive lipase (HSL), is highly active in metabolism of both diglycerides and cholesteryl esters [53]. An alternative means by which lipolysis may influence ERRα/γ activity is via regulation of the ERR coactivators, PGC1α and PGC1β. The ChIP-qPCR results presented here indicate that ERRα is bound to the *Gpr3* promoter even in the “off” state (when the gene is not transcribed). In skeletal muscle, NR corepressor 1 (NCoR1) represses ERRα under basal conditions. However, during stimulated conditions, NCoR1 exchanges for PGC1α which promotes transcription [54]. Therefore, it may be the lipolysis-induced exchange or binding of a coactivator that triggers the “opening” of the region and subsequent transcriptional induction of *Gpr3*.

A key remaining question is why *Gpr3* is not cold induced in all adipocytes upon stimulated lipolysis, considering that *Gpr3* is induced in brown and beige adipocytes but not in canonically white adipocytes [17]. This suggests that the lipolysis-sensitive mechanism facilitating *Gpr3* induction in BAT must be uniquely present in highly thermogenic fat depots. Both ERRα/γ and PGC1α fit this category. The ERRs and PGC1α are enriched in BAT over WAT [36–38], and *Ppargc1a* is induced in thermogenic fat during cold exposure [38].

Importantly, the work presented here does not preclude the possibility of lipolysis-dependent elements and enhancers, located further upstream or downstream of the cold-sensitive promoter. Indeed, the differential transcription profiles of *Ucp1* and *Gpr3* in cold exposed AdipoQ-Cre *Esrra/g* KO mice suggest a more complex transcriptional mechanism of BAT ERRα/γ. While ERRα/γ were required for cold induction of *Gpr3*, they were dispensable for *Ucp1/*UCP1 induction. Conversely, basal expression levels of *Ucp1* were reduced in the ERRα/γ ablated mice and unchanged for *Gpr3*.

While ERRα/γ inverse agonism displays benefits in models of diabetes via hepatic engagement [27,55,56], agonists targeting the ERRs in muscle and BAT may additionally improve metabolic outputs. In skeletal muscle, ERRα/γ enhanced oxidative capacity [46,49,57,58], and Mootha *et al.* proposed that ERRα agonism may offset the molecular implications of skeletal muscle insulin resistance in patients living with type 2 diabetes [39]. In BAT, ERRα/γ mediated the adaptive response to adrenergic stimulation [35], and our results suggest that an ERRα/γ agonist may activate *Gpr3* transcription, thus, providing a therapeutic strategy for harnessing GPR3-induced calorie-burning in thermogenic fat. In line with this, administration of the synthetic pan-ERR agonist, SLU-PP-332, reduced obesity and improved insulin sensitivity in mouse models of metabolic syndrome [59]. Exploring the impact of this compound on *Gpr3* transcription and BAT physiology will further illuminate the therapeutic potential of our findings.

## 5. Conclusions

In conclusion, the cold-induced transcription of *Gpr3* in brown fat is dependent on ERRα/γ, which bind to a cold-sensitive promoter of *Gpr3*. Whether this transcriptional axis directly mediates the lipolysis-dependent element of *Gpr3* transcription remains a subject for further investigation.

## 6. Data and materials availability

Data are available upon request. Plasmids and mouse models used in this study will be available upon request.

## 87. Acknowledgements

The authors wish to thank the members of the Gerhart-Hines group and the Kazak lab for valuable discussions and day-to-day technical advice. This project was supported by the Novo Nordisk Foundation (125973 to ZGH; NNF18OC0033444 to the Center for Adipocyte Signaling). This project has received funding from the European Research Council (ERC) under the European Union’s Horizon 2020 Research and Innovation Programme (Starting Grant aCROBAT agreement no. 639382 to ZGH and Consolidator Grant HEAT-UP agreement no. 101088636 to ZGH). RLM is funded by an EMBO Postdoctoral Fellowship (ALTF 653-2023). CS is a recipient of a Canderel Fellowship. VG was supported by a Foundation Grant from the Canadian Institutes of Health Research (CIHR) (FDT-156254) and a Program Project Grant from the Terry Fox Research Institute (PPG-1091). LK is supported by a Canadian Institutes of Health Research (CIHR) project grant (PJT-190219). The Novo Nordisk Foundation Center for Basic Metabolic Research is an independent research centre at the University of Copenhagen, partially funded by an unrestricted donation from the Novo Nordisk Foundation (NNF18CC0034900 and NNF23SA0084103). Figures 2A and 3A were created with BioRender.com.

## 9. Author contributions

Conceptualization, ZGH, OSJ, LK; investigation ZGH, OSJ, RLM, JR, CS, DML, DT, ALB, VG, LK; Writing – original draft, ZGH, OSJ, RLM; Writing – review and editing, ZGH, OSJ, RLM, JR, CS, DML, DT, ALB, VG, LK; Visualization OSJ and RLM.

