## Supplementary Tables for "Cold exposure induces the constitutively active thermogenic receptor, GPR3, via ERRα and ERRγ"

Supplementary table 1

Reagents:

| **Reagent** | **Company** | **Cat#** |
|---|---|---|
| 293T | ATCC | - |
| DMEM | Wisent | 319-005-CL |
| FBS | Multicell | 098150 |
| Penicillin/streptomycin | Multicell | 450-201-EL |
| Carprofen | Zoetis Canada | Rimadyl injectable solution |
| Lidocain | Vetoquinol | Lurocaine |
| Bupivacaine | Sensorcaine | Bupivacaine Injection BP |
| Polyethylenimine (1 mg/ml) | Polysciences | 23966-1 |
| Optimem | Gibco | 31985 |
| AAVanced Concentration Reagent | System Biosciences | AAV100A-1 |
| AAV helper plasmid pDP8 | Plasmid Factory | PF421-180518 |
| Low-fat diet | Envigo | 2920X |
| High-Capacity cDNA Reverse Transcription kit | Applied Biosystems | 4368813 |
| GoTaq qPCR Master Mix | Promega | A6001 |
| cOmplete™ Protease Inhibitor Cocktail | Roche | 11836170001 |
| BCA assay | Pierce | 23225 |
| 4x Laemmli protein sample buffer | Bio-Rad | 161-0747 |
| Clarity | Bio-Rad | 1705060 |
| Dynabeads protein G | Thermo Fisher Scientific | 10009D |
| SYBR Green I Master Mix | Roche | 4887352001 |

Supplementary table 2

Antibodies:

| **Antigen** | **Company** | **Cat#** | **Dilution** |
|---|---|---|---|
| VCL | Cell signaling | 13901 | 1:5000 |
| CKB | Abcam | ab151579 | 1:1000 |
| UCP1 | Abcam | ab10983 | 1:2000 |
| ERRα | Abcam | ab76228 | 1:1000 |
| GFP | Abcam | ab290 | 1:2500 |
| Anti-rabbit | Promega | W401B | 1:10000 |

Supplementary table 3

Primers:

| **Gene** | **Forward primer** | **Reverse primer** |
|---|---|---|
| *36b4* | TCATCCAGCAGGTGTTTGACA | GGCACCGAGGCAACAGTT |
| *Ucp1* | AAGCTGTGCGATGTCCATGT | AAGCCACAAACCCTTTGAAAA |
| *Gpr3* | ATCACCTGAGCAACCGAGAA | AGATGGGGGTGCATTTTACA |
| *Ckb* | GCCTCACTCAGATCGAAACTC | GGCATGTGAGGATGTAGCCC |
| *Gfp* | AAGGGCATCGACTTCAAGG | TGCTTGTCGGCCATGATATAG |
| *Esrra* | CCAGAGGTGGACCCTTTGCCTTTC | CACCAGCAGATGCGACACCAGAG |
| *Esrrg* | CTCCAGCACCATCGTAGAGGATC | GATCTCACATTCATTCGTGGCTG |
| ChIP *Gpr3* primer set 1 | CATAGTAATGAGGCCTCGCGCC | TAGTGTCCCCAAAACCCCCGAC |
| ChIP *Gpr3* primer set 2 | GGGAGGTCACAGAGGATGTGGG | CAGCTCGTTTCTGCCACCGAAG |
| *Esrra* -3722 neg. | TTGGCATTGATATTGGGGGTGGGAGCAACT | GACTTCTTACTTTGACGCTTTCCTCCATCG |

| | | |
|---|---|---|
| *Prox1* - 55772 (ctrl) neg. | CCAAGCACAAATATCTAATCACCCTTTC | CTTCTTGATAGGTTTATGGGTTGGGC |
| *Essra* (WT and conditional alleles) | CCCTGCTTCTGTGCCCTTTGC | CCACCACTGCCCAGCTTCAC |
| *Essrg* (WT and conditional alleles) | GTTTTAAAGGCCCTTGGTGATCTCGC | CTGCAACCCTTGGACTGCCAGAAC |
| *AdipoQ -Cre* | GACATGTTCAGGGATCGCCAGGCG | GACGGAAATCCATCGCTCGACCAG |
